# Role of interneuron subtypes in controlling trial-by-trial output variability in the neocortex

**DOI:** 10.1101/2022.12.06.519329

**Authors:** Lihao Guo, Arvind Kumar

## Abstract

Trial-by-trial variability is a ubiquitous property of neuronal activity in vivo and affects the stimulus response. Computational models have revealed how local network structure and feedforward inputs control the trial-by-trial variability. However, the role of input statistics and different interneuron subtypes in shaping the trial-by-trial variability was less understood. Here we investigated the dynamics of stimulus response in a model of cortical microcircuit with one excitatory and three inhibitory interneuron populations (PV, SST, VIP). We show that the variance ratio of inputs to different neuron populations and input covariances are the main determinants of output trial-by-trial variability. The effect of input covariances is contingent on the input variance ratios. In general, the network shows smaller output trial-by-trial variability in a PV-dominated regime than in an SST-dominated regime. Our work reveals mechanisms by which output trial-by-trial variability can be controlled in a context, state, and task-dependent manner.

## Introduction

Trial-by-trial variability is a ubiquitous feature of cortical activity *in vivo* (Arieli et al., 1996; Churchland et al., 2010). Instead of being just noise, trial-by-trial variability varies (typically gets reduced) during the stimulus presentation (Churchland et al., 2010; Oram, 2011), due to attentional shifts (Kanashiro et al., 2017) or external stimulation (De Luna et al., 2017). The presence and change in trial-by-trial variability are not merely a statistical property of neuronal activity as it is necessary for behavior (Waschke et al., 2021) and affects the stimulus-response (Arieli et al., 1996) and behavioral performance (Arazi et al., 2017; Rowland et al., 2021). Given the noisy inputs, stochastic neurons, random connectivity, and unreliable synapses, trial-by-trial variability is not surprising. However, the modulation of trial-by-trial variability is usually attributed to local network structure. Computational studies have suggested that the modulation of trial-by-trial variability, particularly the stimulus-induced decrease, can be attributed to the recurrent connectivity in the local network (Litwin-Kumar & Doiron, 2012; Deco & Hugues, 2012; Doiron et al., 2016) and spike level correlations in the feedforward input (Bujan et al., 2015). Network models have also suggested that in a network with excitation and inhibition in balance, a reduction in the variability of excitatory neurons may be accompanied by a corresponding increase in the variability of inhibitory neurons (Kanashiro et al., 2017). Thus, 15 excitatory-inhibitory interactions are crucial for the control of variability.

Previous theoretical works to unravel mechanisms underlying the modulation of trial-by-trial variability have focused on the network of a single excitatory (E) and single inhibitory (I) populations (E-I network) (Litwin-Kumar & Doiron, 2012; Deco & Hugues, 2012; Bujan et al., 2015). However, local networks in the brain are composed of multiple interneuron types (Kepecs & Fishell, 2014). These neuron types differ not only in their chemical signature but also in the neuron-type specific connectivity (Jiang et al., 2015). The interneuron diversity renders the local network with rich dynamical and computational properties which have not been observed in the E-I network (Lee et al., 2018; Hertäg & Sprekeler, 2019; Hahn et al., 2022). However, it remains unclear when and how different interneurons contribute to trial-by-trial variability.

In the neocortex, different interneurons receive specific inputs and selectively inhibit pyramidal cells (PCs). For instance, PV-expressing interneurons are mainly driven by feedforward inputs, inhibiting perisomatic regions and basal dendrites of PCs, whereas SST-expressing interneurons are targeted by top-down inputs through VIP cells, inhibiting apical dendrites of PCs (Larkum, 2013). Moreover, each interneuron type can also be the target of specific neuromodulators. Therefore, by characterizing the contribution of different types of interneurons, we may uncover new mechanisms by which the brain can control the response variability in a context, state, and task-dependent.

To understand the role of interneurons in variability control, we investigated using a model neocortical layer 2/3 consisting of one type of excitatory neuron (Exc) and three types of inhibitory interneuron (PV: parvalbumin, SST: somatostatin, and VIP: vasoactive intestinal polypeptide expressing cells). We refer to it as the EPSV network. First, we characterized the transfer-function of neurons in the EPSV network. Next, we measured trial-by-trial variability at different operative points. In particular, we focused on the input statistics: variance and covariance of input rate to different neuron types. We show that the main determinants of the trial-by-trial variability are the (1) variance ratio of inputs to the different neuron populations and (2) covariance of the inputs. The effect of input covariance is contingent on the ratio of variances. The effect of these variables strongly depends on whether the network is operating in an SST or PV-dominated regime. In general, network connectivity provides a landscape on which input variability could be transformed into output variability by varying the neuron excitability, input connection strengths, and stimulus properties.

## Results

To characterize the role of interneurons in the control of trial-by-trial variability, we study how the trial-by-trial input variability is transferred to the output in a model of neocortex (the EPSV model, see Methods). To this end, we first estimated the transfer-function of a typical neuron in the network operating in an asynchronous-irregular and non-oscillatory state.

### Neuron transfer-function

Experimentally neuron transfer-function is typically estimated by injecting direct current at different amplitudes. However, it is more natural to inject spiking inputs. The number of different types of inputs a neuron may receive depends on the number of neuron types in the network. In a simple scenario consider a network with only one type of excitatory neuron with weak connectivity such that the network remains stable and asynchronous. We injected Poisson-type spiking input at different rates into the network and measured the output firing rates. As expected, in this setting, the output firing rate varies monotonically as a function of the input rate (Figure 1A). If we had injected inhibitory inputs the firing rate would have monotonically decreased from some baseline firing.

**Figure 1:**
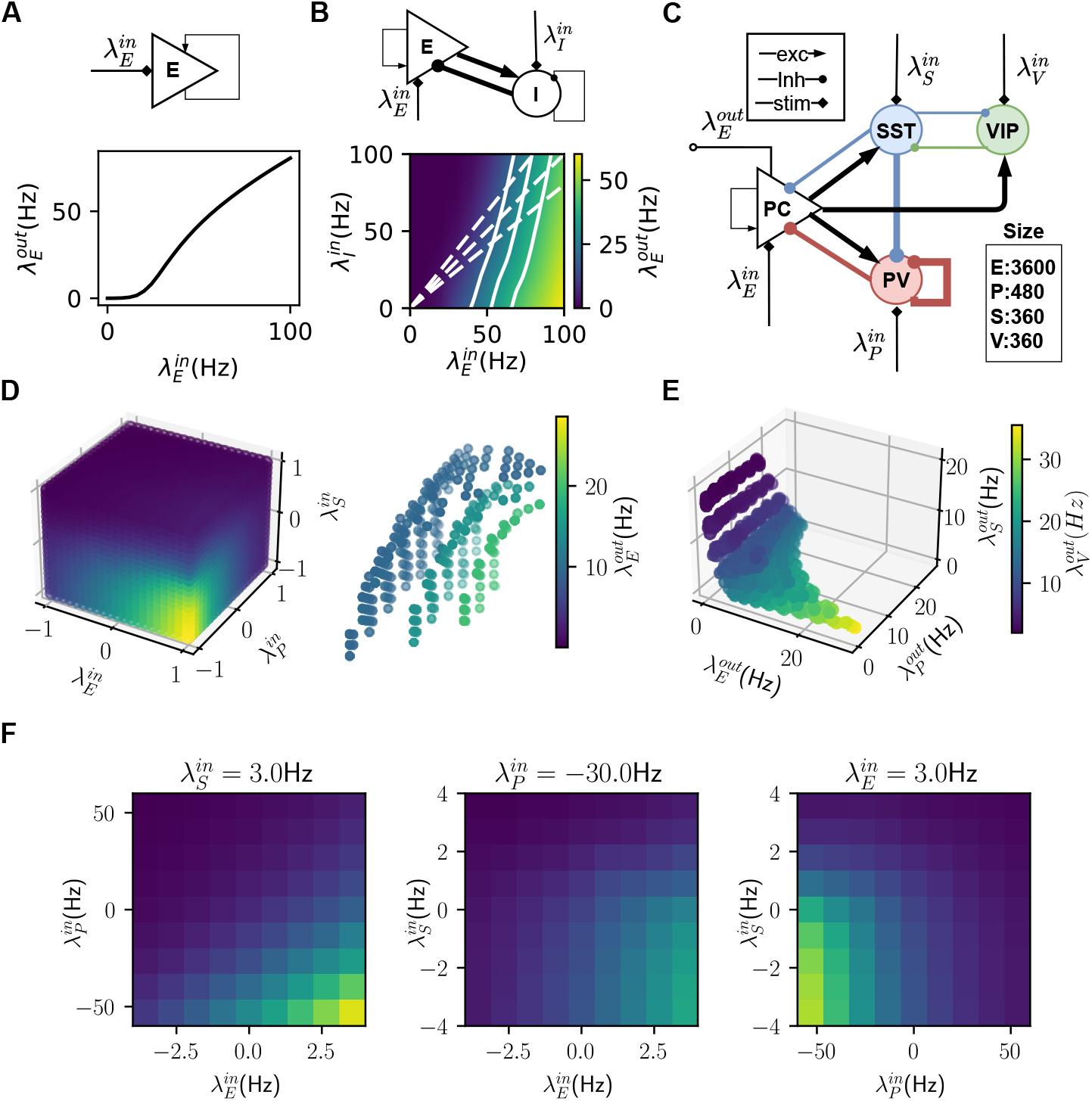
Steady-state neuron transfer-function in different networks. (**A**: Top) Schematic of a one-population network (purely excitatory with weak recurrent connections). (**A**: Bottom) Neuron transfer-function of the one population network where 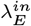 denotes the rates of input (homogeneous Poisson spike trains) to excitatory neurons. (**B**: Top) Schematic of a two-population, excitatory-inhibitory (E-I), network. (**B**: Bottom). Neuron transfer-function of the E-I network (solid white lines show constant output firing (iso-firing) rate contours and dotted white lines show inputs with different E-I ratios). (**C**) Schematic of the EPSV network. (**D**: Left) transfer-function of excitatory neurons in an EPSV network (normalized input). (**D**: Right) Three iso-firing rate surfaces with different output rates. (**E**) Output space of EPSV network with cubic input space as in **D**. The three-dimensional scatter plot shows the activity of E, PV, and SST neurons. The dot color indicated the firing rate of VIP neurons. (**F**) 2-D slices of the transfer-function in **D** to indicate how neuron firing rate changes a function of the input to two populations while the input to the third is fixed. A negative input rate denotes a reduction of input compared to the baseline (see Supplementary Figure S1).

However, when a neuron is a part of a network with excitatory and inhibitory neurons (E-I network) it is important to consider both excitatory and inhibitory inputs separately to determine the neuron transfer-function (Kuhn et al., 2004). This results in a two-dimensional neuron transfer-function (Figure 1B) and a neuron can show identical output firing rate (Exc population) for many different combinations of inputs to Exc and Inh populations (shown as iso-firing rate contours: Figure 1B solid white lines). The neuron transfer-function also reveals how a neuron may respond if the input is varied while maintaining the excitation and inhibition in balance (Figure 1B dotted lines).

Extending the two-dimensional neuron transfer-function to a neuron in the EPSV circuit (Figure 1C), we need to consider the combination of four inputs to the four populations in four-dimensional input space, 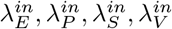 (see section Stimulus evoked input). Because the VIP population and SST population are strongly mutually coupled to each other, we simplified by keeping 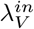 fixed while only changing 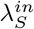 as in (Hertäg & Sprekeler, 2019). We estimated this three-dimensional neuron transfer-function by systematically varying the inputs (see section Stimulus evoked input) to the Exc, PV, and SST populations (Figure 1D left). The negative and positive inputs are relative to the baseline inputs (see section Baseline input). The baseline inputs were chosen to match the network activity to the *in vivo* firing rates of the four neuron types (Yu et al., 2019). Rate traces and raster plots of the network working in baseline state is shown in Supplementary Figure S1 A.

As a function of the input to Exc and PV neurons, the neuron transfer-function was similar to the two-dimensional transfer-function (compare Figure 1B bottom and F left). However, an increase in SST inputs resulted in a threshold-like transition in the output of the Exc population (Figure 1F middle and right panels). This is because of mutual inhibitory coupling between VIP and SST neurons. Increasing the firing rate of SST neurons reduced the activity of VIP neurons which further amplifies SST neurons’ firing rate and effectively silenced the PV neurons. Therefore, at some level of positive input, the network made a transition to the SST-dominated regime characterized by a sharp reduction in the activity of Exc neurons (Figure 1F middle and right panels).

For the three-dimensional neuron transfer-function, we can also define the manifold on which a neuron’s output remains constant despite a change in the input (Figure 1 D right). Next, we rendered the output firing rate of all populations together to visualize all possible network states for a range of inputs to these populations. As expected the recurrent connectivity in the network restricted the possible network states(Figure 1 E). More importantly, this figure shows that when E neurons are firing at high rates, either SST or PV neurons are completely silent. This is a consequence of the switching dynamics of the network which has also been reported earlier (Hertäg & Sprekeler, 2019; Hahn et al., 2022). As we show in the next section, the restricted state space, the shape of iso-firing rate surfaces, and the gradient of the neuron transfer-function play a major role in determining how trial-by-trial input variability is transformed into trial-by-trial output variability.

### Trial-by-trial variability: E-I network

No matter how well we control the experimental conditions, the input to a network always has some trial-by-trial variability. How this input variability is transformed into output variance depends on the gain of the neuron transfer-function in the network. If we consider a single neuron population (Figure 2 A), the task-related input follows a one-dimensional normal distribution (for simplicity) characterized by the mean *μ^in^* and variance *σ^in^* of the input rates across trials. For this case, output variance depends on both *μ^in^, σ^in^*, and the shape of the neuron transfer-function (Figure 2 A,B).

Distinct from the one-dimensional case, there is more freedom in controlling the trial-by-trial output variability in an E-I network. We can visualize the trial-by-trial input variability as a point cloud in a two-dimensional space spanned by inputs to the Exc and Inh populations (Figure 2 B). Each point indicates the input level to Exc and Inh populations in a given trial. It becomes apparent that besides the mean and the variance of the input to Exc and Inh neurons, the shape and orientation of the input point cloud are also important to determine the output variance. When the input point cloud is elongated (elliptical) and aligned to the iso-firing rate lines, output variance (contributed only by Poissonian fluctuation) will be small, irrespective of the individual input variances.

**Figure 2:**
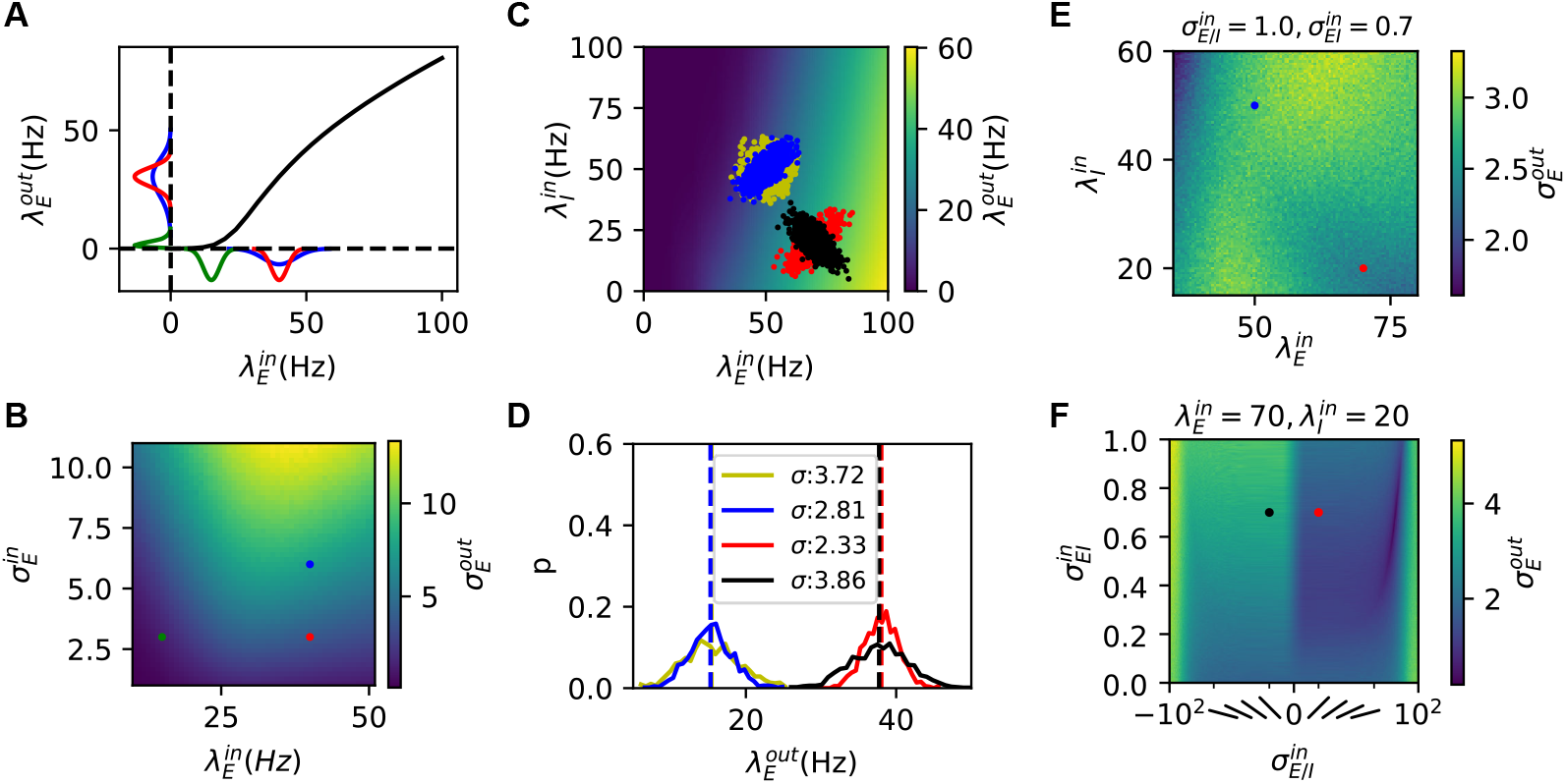
Transformation of trial-by-trial input variability to output variability. (**A**) A normal distribution of inputs across trials (x-axis) is transformed into a distorted distribution of outputs (y-axis). Different colors denote different distributions. (**B**) Output trial-by-trial variability (measured as standard deviation) for different mean and variance across-trial inputs in a one-dimensional case. (**C**) Transfer-function of the E-I network with weak recurrency. The four input clouds (denoted by colors) were sampled from distinct trial-by-trial input distributions. (**D**) Output rate distribution of corresponding input clouds shown in **C** for the E-I network. (**E**) Output trial-by-trial variability as a function of mean input to Exc and Inh populations for a fixed covariance matrix. The red(blue) dot denotes the red(blue) distribution in **C**. (**F**) Output trial-by-trial variability as a function of covariance 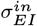 and variance ratio 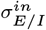 (slated bars illustrate the orientation of sampling cloud), for a fixed mean input to the two populations. The red(black) dot denotes the red(black) distribution in **C**.

It suggests that, for the two-dimensional case, we need to consider the complete statistics of the inputs to the two populations. For simplicity, we assumed that the inputs across trials follow a bivariate normal distribution. In this setting, we need to define the following to characterize the output variance:

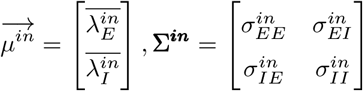

We decomposed the distribution of the input point cloud into three factors: the trial-by-trial covariance 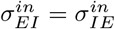 (compare blue and yellow points in Figure 2 C), trial-by-trial EI balance 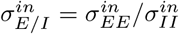 (compare red and black in Figure 2 C) and the input mean 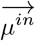 (compare blue and red Figure 2 C). In addition, we assumed that the total input variance was fixed 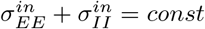.

First, we fixed the trial-by-trial covariance and variance of inputs to Exc, and Inh populations and systematically varied the Exc, and Inh input means. We used the neuron transfer-function, estimated from network response during ongoing activity state, to obtain the corresponding output rates. The trial-by-trial variability reflected the gradient of the 2-dimensional neuron transfer-function and varied greatly as we change the input means (Figure 2 E). Typically, low firing rate regions were associated with high variability (Figure 2 E). The results in Figure 2 E show that in an E-I network if the input covariance matrix remains fixed, just a change in the mean input, which drives the network to a higher firing rate, is sufficient to reduce trial-by-trial variability (Figure 2 C blue and red histogram, and Figure 2 E blue and red dots) unless we are operating at very low evoked firing rates. This observation provides a new explanation of the reduction in the trial-by-trial variability during evoked activity (Churchland et al., 2010). However, from the Figure 2 E we can also derive constraints on Exc and Inh inputs which will increase trial-by-trial variability during evoked activity.

Next, we systematically varied covariance between inputs to Exc and Inh neurons and the variance ratio of inputs to Exc and Inh neurons while keeping the mean of inputs constant (Figure 2 F). This analysis is consistent with the idea that output variance is minimal when the input point cloud is elliptical with a positive slope and aligned to the iso-firing rate lines. It shows that when Exc and Inh inputs are anti-correlated (Figure 2 black distribution), the input point cloud is orthogonal to the iso-firing rate lines resulting in maximal output variance.

### Trial-by-trial variability: EPSV network

Next, we used the same approach described in the previous section to isolate the role of different interneurons in shaping trial-by-trial variability. First, we fixed the covariances and variances (assuming a standard normal distribution) and systematically varied the mean of the input to different neurons. Given that network interactions render the neuron transfer-function nonlinear, the trial-by-trial variance was also highly nonlinear as a function of mean input (Figure 3 A left). Unlike the standard E-I network (Figure 2 E), in the EPSV network for most cases, output variability was positively correlated with output rate (Figure 3 A right). However, it was possible to find input configurations when an increase in output firing rate was associated with a decrease in trial-by-trial variability (as is experimentally observed (Churchland et al., 2010)). But such a reduction was only possible for high firing rates when neurons’ transfer-function saturated (Figure 3 A black dots).

**Figure 3:**
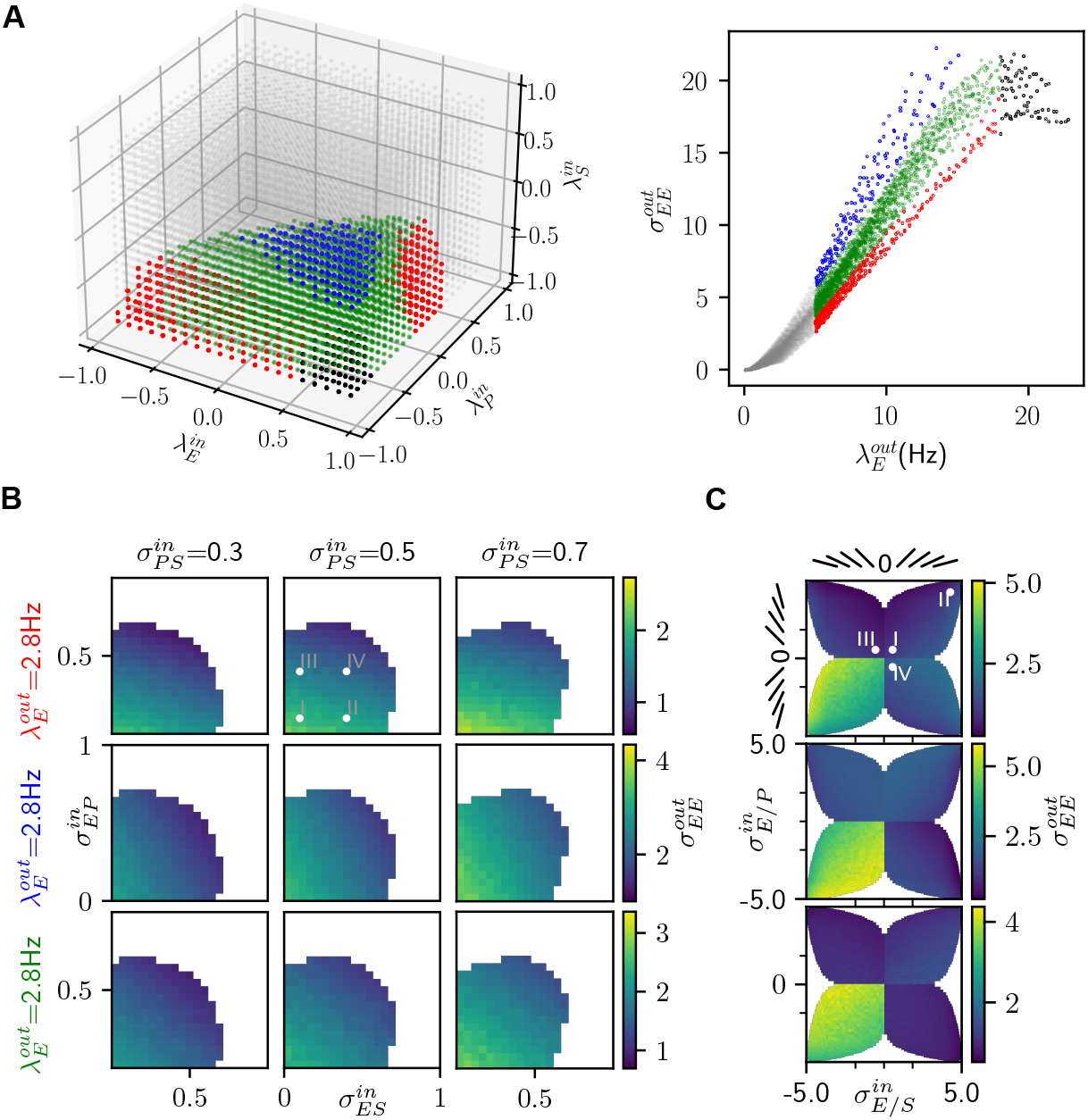
Trial-by-trial variability in EPSV circuit derived from neuron transfer-function. (**A left**) Output trial-by-trial variability (quantified as variance) of E population as a function of across-trial input means with a fixed covariance matrix (inputs were sampled from an un-correlated tri-variate normal distribution). The color code is in the right panel. (**A right**) Output trial-by-trial variability as a function of output rate of E neurons. Each dot corresponds to a specific input (see the left panel). Red, blue, and green dots correspond to the network in a PV-dominated, SST-dominated, and both influencing regime. Black dots refer to a high firing rate region when output variance decreases when increasing the output firing rate. (**B**) Output trial-by-trial variability as a function of the trial-by-trial covariance of inputs with a fixed variance 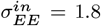, 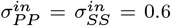. From the top row to the bottom row, the network was tuned to operate in PV-dominated (red) or SST-dominated (blue), or PV-SST driven (green) regimes with similar output rates. (see Supplementary Figure S1 A-C for spiking activity rasters.) (**C**) Output trial-by-trial variability as a function of the trial-by-trial variance ratio with fixed covariance 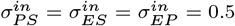. Slanted bars indicate the orientation of the input point cloud. Roman numerals in the **B** and **C** refer to the four cases simulated in Figure 5 and Figure 4.

We also noted that just based on the gradient of the neuron transfer-function, trial-by-trial variability could be roughly divided into three regimes (Figure 3 A right). Above 5 Hz output rate, the slope of output firing rate vs trial-by-trial variability curve was smallest when the network was operating in a PV-dominated regime (Figure 3A red dots) and the slope was largest in an SST-dominated regime (Figure 3A blue dots). Below 5Hz firing rate (Figure 3 A gray dots), we investigated the effect of the trial-by-trial input covariance matrix on the trial-by-trial output variability. We chose a low firing rate region because in this regime all neuron populations were spiking and influenced the output variability. For the high firing rate region, the system degenerated into a lower dimension, e.g. reverting to the E-I network (Figure 2 C-F), when either PV or SST neurons were almost silent (Supplementary Figure S2 A). However, the results of trial-by-trial variability transfer were similar for both regions (Figure 3 and Supplementary Figure S2).

Next, we varied the trial-by-trial input variance ratio or trial-by-trial covariance while keeping the mean of inputs constant (Figure 3 B,C). To make a fair comparison, we chose three different mean input configurations such that the network worked in either PV-dominated (Figure 3 B,C top) or SST-dominated (Figure 3 B,C middle) or both controlling (Figure 3 B,C bottom) regime while having approximately the same output firing rate (E population). Spiking activity and firing rate for the three operating regimes are shown in Supplementary Figure S1 A-C.

The input variance ratio (orientation of the input point cloud) played a major role in controlling the trial-by-trial output variance of the excitatory population (Figure 3 C). To quantify the effect, we varied the ratio of input variances while keeping the covariance fixed such that the input point cloud ranged from completely aligned to orthogonal to the iso-firing rate manifolds (Figure 3 C) similar to the 2-dimensional case (Figure 2 C,F). In the PV-dominated regime (Figure 3 C top), the orientation for input point cloud with the lowest trial-by-trial output variability was in the region where inputs to E-P pair 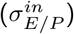 were positively aligned and to E-S pair 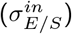 negatively aligned. Consequently, only inverting the sign of trial-by-trial input covariance of the E-P pair from positive to negative induced a large change of trial-by-trial variability (Figure 3 C top, see also Figure 2 F for 2-D case). As expected, the situation reversed for the SST-dominated regime (Figure 3C middle). Finally, both 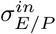 and 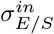 contributed to the output variance with comparable strength in the regime where both PV and SST interneurons shaped the network response (Figure 3 C bottom). For all operating points, the lowest trial-by-trial output variability was in the region of correlated trial-by-trial inputs to both E-I pairs (input cloud with positive slope), whereas the highest variability was in the region of anti-correlated trial-by-trial inputs to both E-I pairs (input cloud with negative slope).

To analyze the effect of covariance, we assumed a fixed ratio of trial-by-trial input variances to the different neuron populations where the input point cloud was in the region of correlated inputs to both E-P, E-S pairs (Figure 3C I). Next, we varied the covariance between different pairs of populations (Figure 3 B). In the PV-dominated regime (Figure 3 B top row), increasing 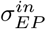 decreased the trial-by-trial variance, whereas 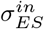 had a much smaller effect. In the SST-dominated regime (Figure 3 B middle row) increasing 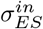 decreased the trial-by-trial variance, whereas 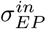 had a much smaller effect. Finally, when both PV and SST neurons affected the network activity, an increase in both 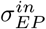 and 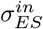 resulted in a decrease in the output trial-by-trial variability (Figure 3 B bottom row). These results show that the covariance between inputs to the three neuron populations affected the trial-by-trial variability in a state-dependent manner.

### Trial-by-trial variability: Network simulation

To confirm the analysis above derived from interpolated neuron transfer-function, we simulated the stimulusresponse of the EPSV network (see Methods section: Trial-by-trial variability of the input). We tuned the EPSV network in a PV-dominated regime and stimulated the three neuron populations with inputs sampled from a multivariate normal distribution with a specific covariance matrix (Figure 4 A top row, and B). In each trial, the network responded with a different output rate (Figure 4C). To quantify the trial-by-trial variability we measured the covariance matrix from the activity of the three populations (Figure 4 A bottom row).

**Figure 4:**
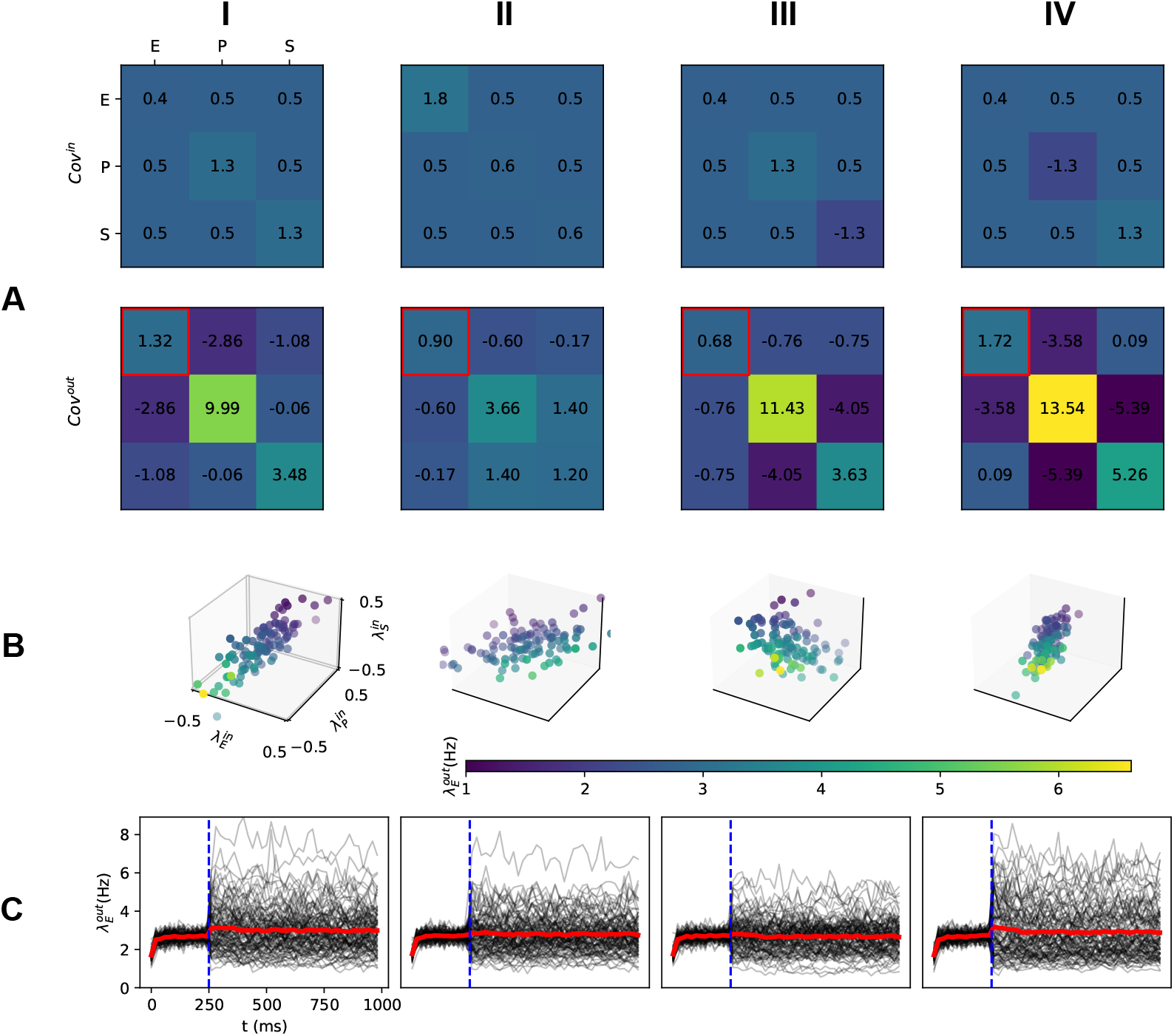
Simulation of trial-by-trial variability controlled by input variance ratio in a PV-dominated regime. (**A top**) The input covariance matrix for sampling input point clouds (**B**). From **I** to **II**, the trial-by-trial input variance ratio between E and SST/PV populations were increased. In column **III** and **IV**, negative variance in *Cov^in^* denotes the negative slope of input point clouds as shown in Figure 2 F. The network operated in a state corresponding to the top row in Figure 3 C. (**A bottom**) Output covariance matrix across 100 trials where inputs for each trial were sampled from the covariance matrix in **A top**. The output variance of the excitatory population is indicated with a red square. (**B**) Trial-by-trial input point clouds sampled from given covariance matrix as columns **I** to **IV** in **A top** (normalized range). **C** Peristimulus time histogram (PSTH) of excitatory population response. Black lines: individual trial. Red line: average response over 100 trials. Stimulus (which involved only a change in the variance and/or covariance without any change in the mean) was provided at 250 ms, and the output covariance matrix in the panel **A bottom** was calculated for the last 500 ms.

First, we simulated the effect of the ratio of trial-by-trial input variance while keeping other input variables fixed (Figure 4 A top row B). We assumed that the total variance remains constant (i.e. the sum of diagonal in Figure 4 A top row). The results are consistent with the estimate of trial-by-trial variance derived from the neuron transfer-function. For instance, changing the ratio of the trial-by-trial input variance to the E and P neurons (by an increase of trial-by-trial input variance to the excitatory population while decreasing the variance of input to the PV population) resulted in a decrease in the output variability (Figure 4 A I, II). In this example, a change in the input variance ratios altered the alignment of the input point cloud with the iso-firing rate surfaces, therefore, we observed a decrease in the trial-by-trial output variance.

To further illustrate the effect of the orientation of the input point cloud (or the ratio of input variances), we simulated the network response when the input point cloud had a negative slope (Figure 4B III, IV). In the PV-dominated regime, the slope of the E-S input point cloud affected the output trial-by-trial variability by a small amount (Figure 4 C III). However, when the input cloud slope was negative in the E-P dimensions, trial-by-trial output variability was very high (compare Figure 4 C II and IV), consistent with the results obtained using neuron transfer-function (Figure 3 C top).

Next, we varied the trial-by-trial covariance between the inputs to the three neuron populations while fixing all other variables (Figure 5 A top row). The effect of input covariances is best seen when the input cloud is aligned to iso-firing rate surfaces. Therefore, we tuned the input variance ratios accordingly (see diagonal in Figure 5 A top row and Figure 3 C II). With these variance settings, an increase in the covariance between inputs to E and S populations resulted in a small decrease in the output variance of the excitatory population (red square in Figure 5 A bottom row). Because we had tuned our network in a PV-dominated regime, an increase in the covariance between inputs to E and P populations resulted in a much larger decrease in the trial-by-trial variability (compare columns I and III in Figure 5A top row) resulted in a much larger decrease in the variance of E population (compare Figure 5 C I and III columns). Finally, when we increased both the covariance (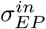 and 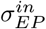), we also observed a large decrease in the trial-by-trial variability of E neurons (compare Figure 5 I and IV columns). In general, consistent with our estimates from the neuron transfer-function, in numerical simulations of evoked responses we found that indeed, trial-by-trial variability can be controlled by varying the variance ratio and covariance of trial-by-trial inputs to the different neuron types.

**Figure 5:**
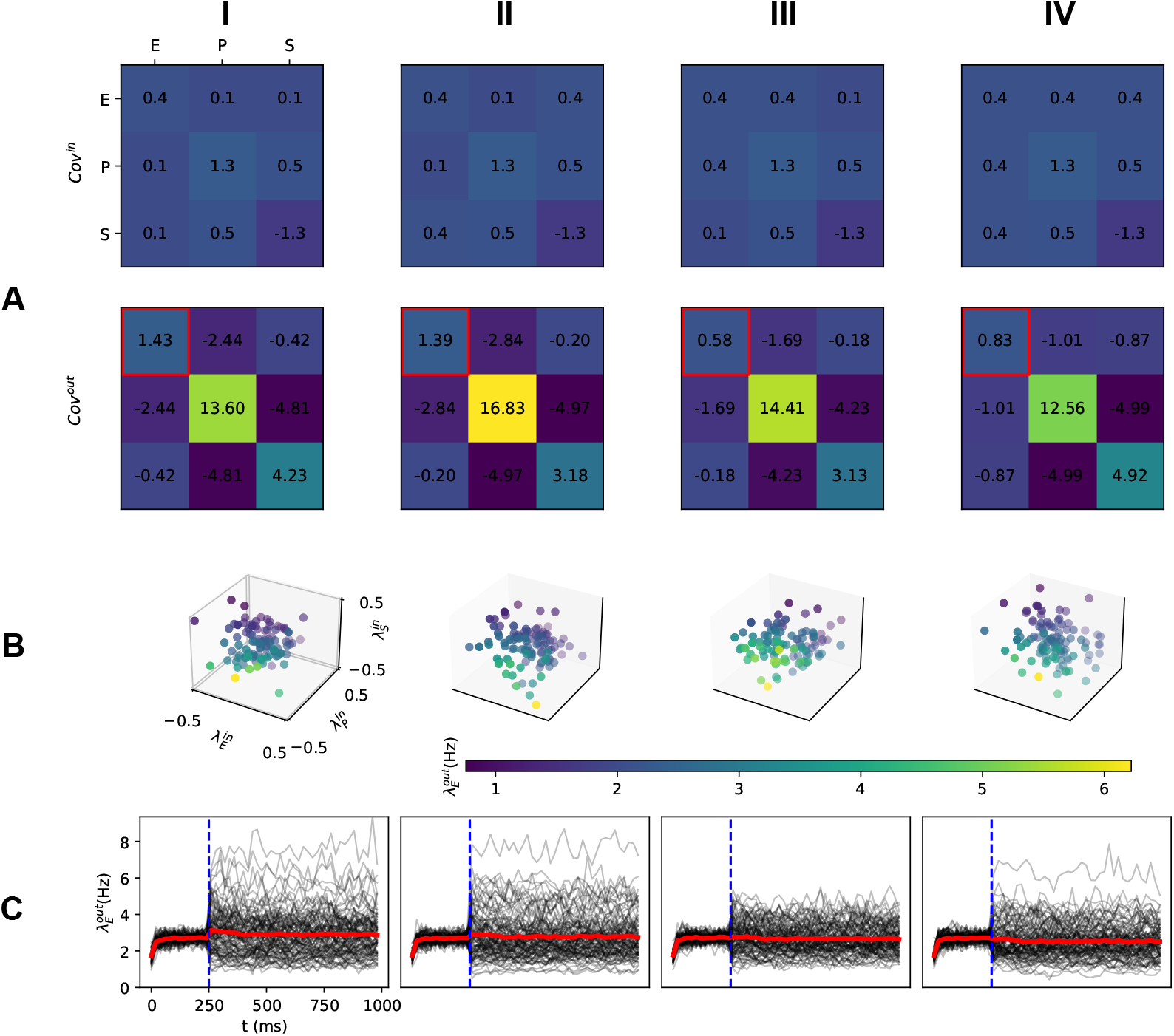
Simulation of trial-by-trial variability controlled by input covariance in a PV-dominated regime. (**A,B,C**) Same arrangement as in Figure 4. From column **I** to **II** (**III**), trial-by-trial input covariance between E and SST (PV) populations was increased. In column **IV**, covariances between E-P and E-S pairs were both increased.

Note that Figure 5 IV had a larger output variance compared to Figure 5 I which is not exactly the same with Figure 3 B middle panel top row. This is due to linear interpolation of neuron transfer-function and limited sampling size for simulations while the actual underlying iso-firing rate surfaces are nonlinear Figure 1 D right. Simulations of trial-by-trial variability in SST-dominated regime (Supplementary Figure S4) were also consistent with the estimations made from neuron transfer-function (Figure 3 B,C middle row). For high firing rate region, simulations of trial-by-trial variability in PV-dominated regime (Supplementary Figure S3) had similar result to low firiging rate region (Figure 5 and Figure 4) which were consistent with estimations made from neuron transfer-function (Supplementary Figure S2 B,C top row).

### Trial-by-trial variability of inhibitory neurons

In our model, the change of trial-by-trial output variability accompanied different correlations between E, P, and S populations (Figure 5 A bottom and Figure 4 A bottom). Increasing the trial-by-trial input variance to the excitatory population does not necessarily cause a corresponding increase in trial-by-trial output variance (compare Figure 4 A columns I and II). Instead, the outputs of different populations decoupled (compare the output covariance between E and P/S populations in Figure 4A columns I and II). In the PV-dominated regime, the trial-by-trial output variability (E population) was mainly controlled by the high-firing level PV neurons. Therefore, an increase or decrease of output variance accompanied a larger or smaller anti-correlation between the outputs of the E-P pair (compare columns in Figure 5). The network connectivity generated negative correlations between the outputs of E-I pairs across trials. Such a negative correlation could be enhanced or canceled by the distribution of inputs across trials. Consequently, the interneurons (PV and SST neurons) contributed more or less to output variance (E population).

Whether the variability of inhibitory interneurons changes in the same way as the variability of the excitatory population depends on the network connectivity. In our model, the output variability of inhibitory populations depends on how their iso-firing rate manifold is aligned with that of the excitatory population. To illustrate this, we considered a two-population E-I network. This network can operate in two regimes – weak self-coupling (or strong mutual interactions) and strong self-coupling (weak mutual interactions). For the weak self-coupling regime, E-I populations had a strong mutual interaction such that the change of one population influenced the other (Figure 6 B left column). In this regime, E and I iso-firing rate lines were not aligned, therefore a decrease in the variability of the excitatory population will be accompanied by an increase in the variability of the inhibitory population. A similar argument has been made by (Kanashiro et al., 2017). By contrast, in the strong self-coupling regime, the output rate of a population did not depend on the input from the other population (Figure 6B right column), and iso-firing rate lines of both E and I populations were aligned. Therefore, a decrease in the variability of the excitatory population will be accompanied by a corresponding decrease in the variability of the inhibitory population.

**Figure 6:**
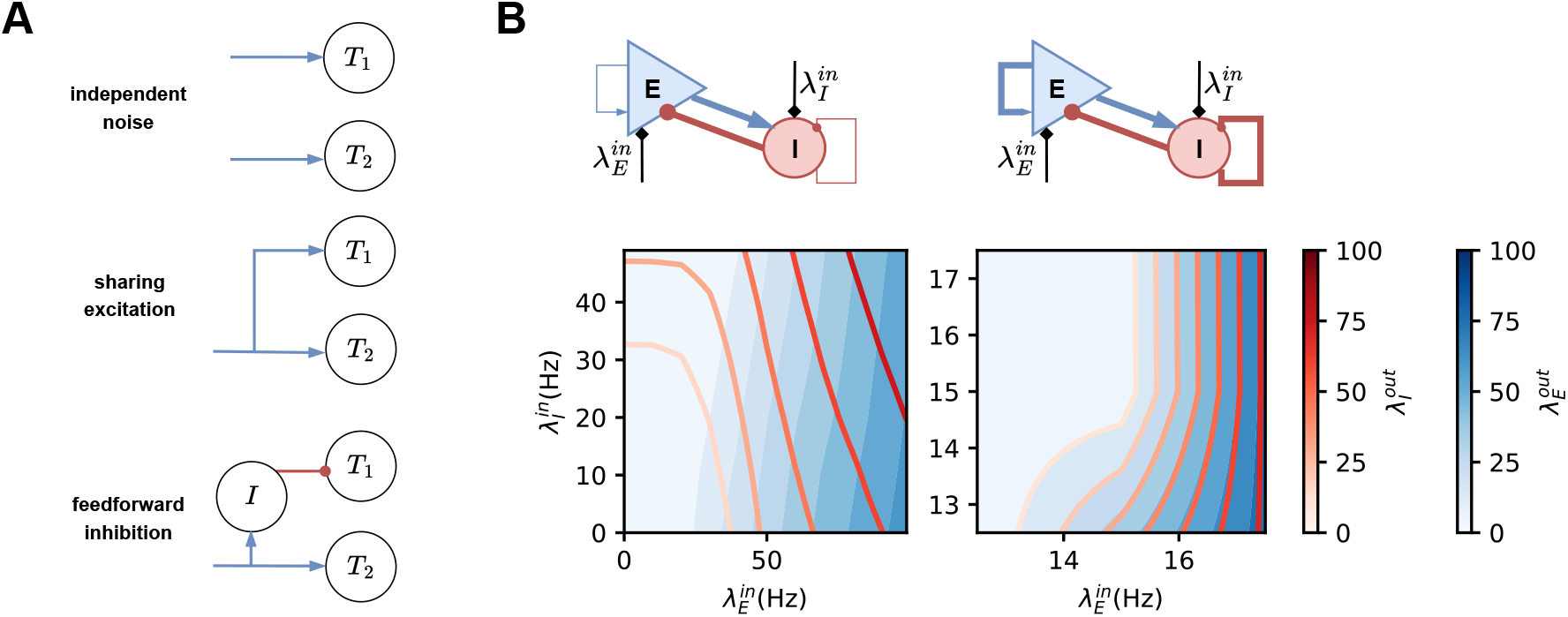
Mechanism of trial-by-trial variability control. (**A**) Trial-by-trial input distribution modulated by feed-forward input connectivity. (**top**) Two target populations receive independent background input. (**middle**) Target populations receive shared excitatory input. (**bottom**) Target populations receive shared input with opposite signs. (**Bleft**) Contours of transfer-function (blue for 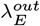 and red for 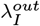) in a weak recurrently connected E-I network. In this configuration iso-firing rate lines between Exc and Inh populations do not align. (**Bright**) In a strong recurrently connected E-I network. In this configuration iso-firing rate lines between Exc and Inh populations align.

## Discussion

Here we have investigated how the mean, variance, and covariances of trial-by-trial inputs affect the trial-by-trial variability of the neocortical activity. Transfer of input distribution to output is mediated by the neuron trasfer-function. The number of distinct neuron types in a network determines the dimensionality of the neuron transfer-function. For the neocortical microcircuit with one excitatory and two or three inhibitory populations, the neuron transfer-function becomes 3- or 4-dimensional. As the dimensionality of the neuron transfer-function increases beyond one, we need to consider not only the variance of the input but also the ratio of input variances and covariance between all pairs of neuron types. Here, we have unraveled how the ratio of variance and covariance of inputs to excitatory, PV, and SST neurons affect the trial-by-trial variance of cortical activity.

For the neocortex network model, the non-linearity of steady state neuron transfer-function resulted in non-trivial relation between output rate and output variance (Figure 3A). Next, the output variance could be significantly altered without any change in the output rate by tuning the input covariance matrix ((Figure 3 B, C). In general, the shape and orientation of input distribution across trials should align with the iso-firing rate manifold for the corresponding network state to reach low output variability. Finally, the non-linearity of the neuron transfer-function implies that the input covariance matrix may never fully align with the iso-firing rate manifolds hence the trial-by-trial output variability is inevitable. The orientation of the input cloud (ratio of variances) is the main predictor of trial-by-trial variability as it aligned the input point cloud to the iso-firing rate manifold. Once such an alignment is achieved, covariance can be varied to further modulate the trial-by-trial variability.

### Control of trial-by-trial variability

Trial-by-trial variability is necessary for behavior (Waschke et al., 2021) therefore it is crucial that it can be varied in a context-, behavioral state-, and task-dependent manner. For instance, during the early stages of learning, we would like higher variability to explore the state space but once the task is learned animals should reduce variability in their behavior.

Our model suggests that feedforward connectivity provides a natural way to alter the input statistics. If the two target populations receive independent inputs (Figure 6 A top), there is no correlation between inputs, and the output variability is solely controlled by the trial-by-trial input variance. In such a scenario, feedforward weights can modulate the input variance: the larger the feedforward weights, the larger the trial-by-trial input variance, and vice versa. When the target populations, e.g. E and P populations, share a certain level of input excitation (Figure 6 A middle) due to feedforward divergent connections from the same sources (thalamus), their inputs co-vary and the slope of the input point cloud is positive. In this scenario, the degree of shared input controls the degree of covariance (Figure 3 B and Figure 5), and feedforward input strength controls the input variance ratio i.e. the slope of the input point cloud (Figure 3 C and Figure 4 column I, II). The slope of the input cloud could be inverted if the common input to one of the two populations is mediated via an inhibitory interneuron (Figure 6 A bottom). Thus, the structure and strengths of feedforward connectivity provide a natural control over the distribution of received inputs across trials. Feedforward input synapses can be learned through plasticity mechanisms so that the animal may reach the desired degree of trial-by-trial variability in their activity and thereby in their behavior. Furthermore, modulation of neuronal excitability and synaptic strength by neuromodulators can provide a context-dependent control of the trial-by-trial variability.

In parallel to the feedforward input structure, changes in the local activity dynamics can also modulate trial-by-trial variability by changing the gradient of the neuron transfer-function and the curvature of iso-firing rate curves. At the simplest, this can be achieved by switching the network between SST-dominated and PV-dominated states. Furthermore, intrinsic excitability and synaptic plasticity, when changing the network between weak (Figure 6 B left) and strong connection (Figure 6 B right) regimes, can modulate trial-by-trial variability by altering the whole landscape of neuron transfer-function.

### Relationship with other explanations of trial-by-trial variability control

Previous models of trial-by-trial variability have mostly focused on the dynamics of recurrent connectivity (Litwin-Kumar & Doiron, 2012; Deco & Hugues, 2012; Doiron et al., 2016; Schnepel et al., 2015). However, trial-by-trial variability is also shaped by external inputs (Oram, 2011; White et al., 2012) and attention signals (Kanashiro et al., 2017; Doiron et al., 2016). More specifically, within-trial correlations in the feedforward inputs can modulate the trial-by-trial variability (Bujan et al., 2015).

Here we have further explored the role of feedforward inputs in shaping the trial-by-trial variability in a network with three different interneuron populations. While focusing on the feedforward input rate covariance and variances, we have not ignored the role of recurrent activity dynamics. The neuron transfer-function responsible for the transfer of input variance and covariances is shaped by the recurrent activity. That is, the neuron transfer-function we have used is not the same as we may estimate by current injections in a silent network (e.g. *in vitro* slices). When the network is operating in different regimes such as an inhibition stabilized network (ISN), the results may change but only to the extent that in an ISN regime neuron transfer may be quite different (Sanzeni et al., 2020). Thus, the approach of using the neuron tranfer-function allows us to combine both input statistics and recurrent activity state.

Thus, in contrast to previous work where the focus was on the input correlation at the spiking activity level, we showed that interneurons contribute to the rate variability in three ways by modulating: (1) the effective transfer-function of the neuron, (2) the operating regime of the network and (3) the trial-by-trial input variance and covariances.

### Function of interneurons

Given the diversity of interneurons in the brain, there is an impetus to identify specific functions of each interneuron subtype (Kepecs & Fishell, 2014). Classically, the function of interneurons is thought in terms of control of gain modulation (Silver, 2010; Isaacson & Scanziani, 2011), control of network activity state (Brunel, 2000), decorrelation of network activity (Renart et al., 2010; Tetzlaff et al., 2012), gating of input (Isaacson & Scanziani, 2011; Kremkow et al., 2010) and control of dynamics of oscillations (Lee et al., 2018; Hahn et al., 2022). Moreover, different interneurons such as PV, SST, and VIP have been implicated in specific functions such as layer-specific control of excitation-inhibition balance (Naka et al., 2019) and synchronization of gamma-band oscillations (Veit et al., 2017). Here we have identified a new role of interneurons – in controlling the trial-by-trial variability. In particular, for the first time, we highlight the importance of the structure of task-related feedforward inputs to the interneurons. Moreover, multiple interneuron types provide more means to control the trial-by-trial variability. Because the cortical networks can switch between SST and PV-dominated regimes depending on the context and attention levels of the animal, interneurons provide mechanisms to modulate trial-by-trial variability in a context-, task-, and behavioral state-dependent manner.

### Model limitations

In the model, we have a number of assumptions e.g. all neurons were modeled as point neurons and connectivity was independent of spatial distance among neurons. However, the most crucial assumption we made is that the network is operating in a near asynchronous-irregular state. In such a state, it is straightforward to relate the input variance and covariance to output variance. But when the network is operating in an oscillatory state (Supplementary Figure S1 D), our single neuron transfer-function approach will not be sufficient. Given the oscillations, we will also have to consider the serial correlations in the input spiking activity and input oscillations. Moreover, inputs with serial correlations or oscillations could affect the downstream network by entrainment or resonance (Hahn et al., 2014). While interesting, this is beyond the scope of the current manuscript.

Next, our conclusion that input covariance and variance ratio are crucial for the control of trial-by-trial variability is contingent on the fact that the iso-firing rate surfaces are continuous. It is possible that due to non-linear interactions, iso-firing rate regions in the input space appear as small islands. In such a scenario, the effect of input variance and co-variance also become contingent on the operating point or the mean output firing rate.

We have shown that increasing the covariance of inputs to E and PV or SST neurons reduces the trial-by-trial variability. This is true when input variance is small enough such that the input cloud can align with the iso-firing rate manifold. When input variance becomes larger, or the curvature of iso-firing rate surfaces is high, an increase in the covariance can misalign the input point cloud from the iso-firing rate cloud and result in a higher trial-by-trial variability. Moreover, when local activity is high, it might end up altering the input statistics and render our predictions wrong.

### Model predictions and model verification

Like any good computational model, we have also made several simplifications in our model (see Methods). However, our model still captures a number of crucial biological details and makes testable predictions. First and foremost, our results suggest that to better understand the modulation of trial-by-trial variability we should measure the variability and co-variability of inputs to different neuron populations. A straightforward prediction of our model is that when inputs to excitatory and inhibitory neurons are correlated, an increase in the input variance may not increase the output trial-by-trial variability. Recently, it has become possible to elicit behavior by optical stimulation of selected neurons which were also activated by an external stimulus (Marshel et al., 2019). Our model predicts that under optogenetic activation/inactivation the trial-by-trial output variability should be different from that observed during sensory stimulation conditions because input structures are qualitatively different in the two experiments.

Our model also predicts that there will be higher trial-by-trial variability when the network is operating in an SST-driven state. Interestingly, a reduction in SST activity is often seen in experiments, e.g. non-whisking and whisking in the barrel cortex of mice (Yu et al., 2019). The feedforward inhibition circuit motif ensures that excitatory and inhibitory inputs to a neuron are correlated. This kind of correlation in our model would lead to low trial-by-trial variability. So we predict that blocking of feedforward inhibitory motif should not only alter the response mean firing rate but also increase the trial-by-trial variability.

Neuromodulators can alter synaptic strengths and neuronal excitability. In our model, a change in the feedforward synaptic weights implies a change in the input variance and a change in neuronal excitability implies rotation or shift of the neuron iso-firing rate manifolds. Therefore, the effect of any neuromodulator on trial-by-trial variability should be non-monotonic because only for a specific amount of neuromodulator it may be possible to align the input cloud with the iso-firing rate manifold.

## Methods and Materials

### Neuron model

The neurons were realized as the adaptive exponential integrate and firing model (Brette & Gerstner, 2005):

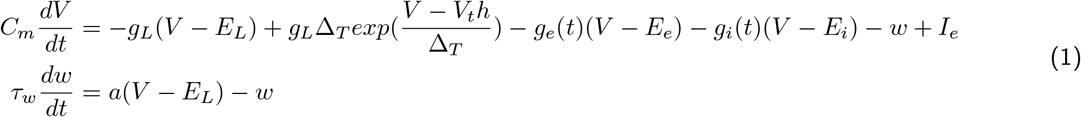

Neuron parameters are provided in the Table 1 (Lee et al., 2018).

**Table 1:**
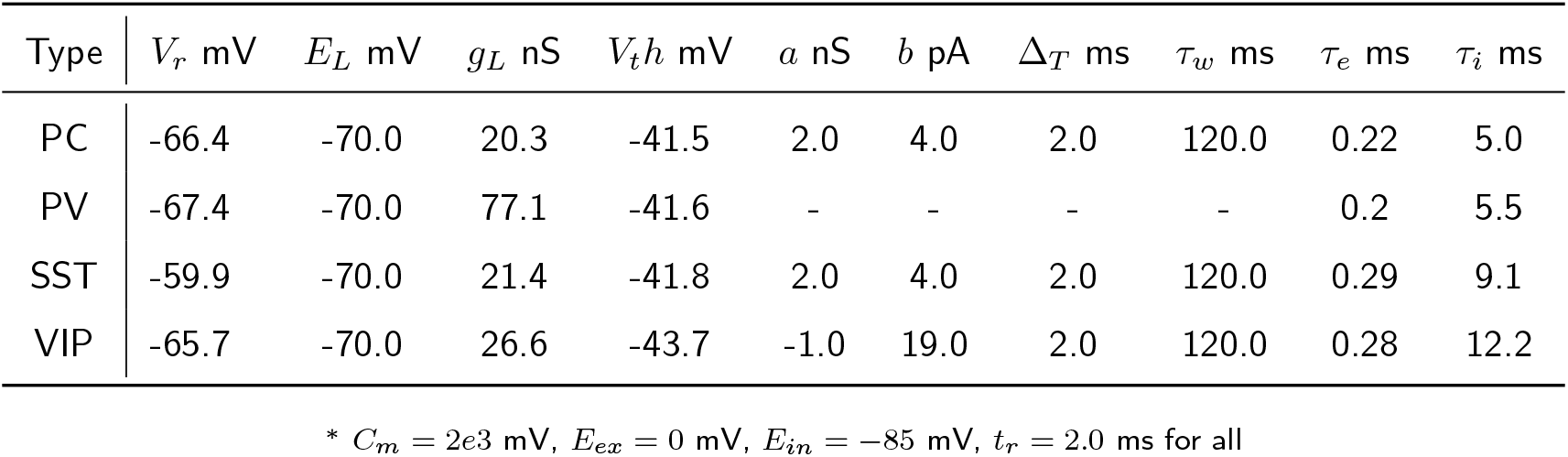
Neuron parameters*

### Synapse model

Neurons were connected using conductance-based synapses. Each incoming spike resulted in a conductance transient which decayed exponentially with a time constant *τ_syn_*, e.g. *τ_e_* for excitatory synapses and *τ_i_* for inhibitory synapses:

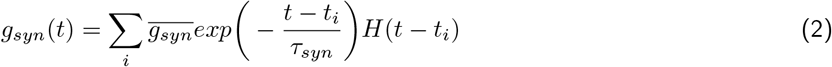

where *t*_i_ is the arrival time of *i^th^* spike and *H* is the heaviside step function.

### Network model

The network consists of 4800 neurons with 3600 excitatory, 480 PV, 360 SST, and 360 VIP neurons (Rudy et al., 2011). Neuronal connectivity parameters (see Table 2) were taken from (Hertäg & Sprekeler, 2019) with modifications. Given the size of the network, neurons were connected in a distance-independent manner that each pair of neurons has probability *p_ts_* to form a connection depending on the types of source and target as in Table 2 left. For synaptic conductance, we first chose the value 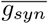 for each pair of connections as in Table 2 right. Next, we scaled each synaptic weight by a number randomly drawn from a log-normal distribution (*μ* = 0, *σ* = 1) and upper bounded the weights by 0.5 mV (−2.0 mV) for EPSP (IPSP). The distribution of excitatory conductance is shown in Figure 7. In the main text, all the results are shown for a network with log-normal synaptic weight distribution.

**Figure 7:**
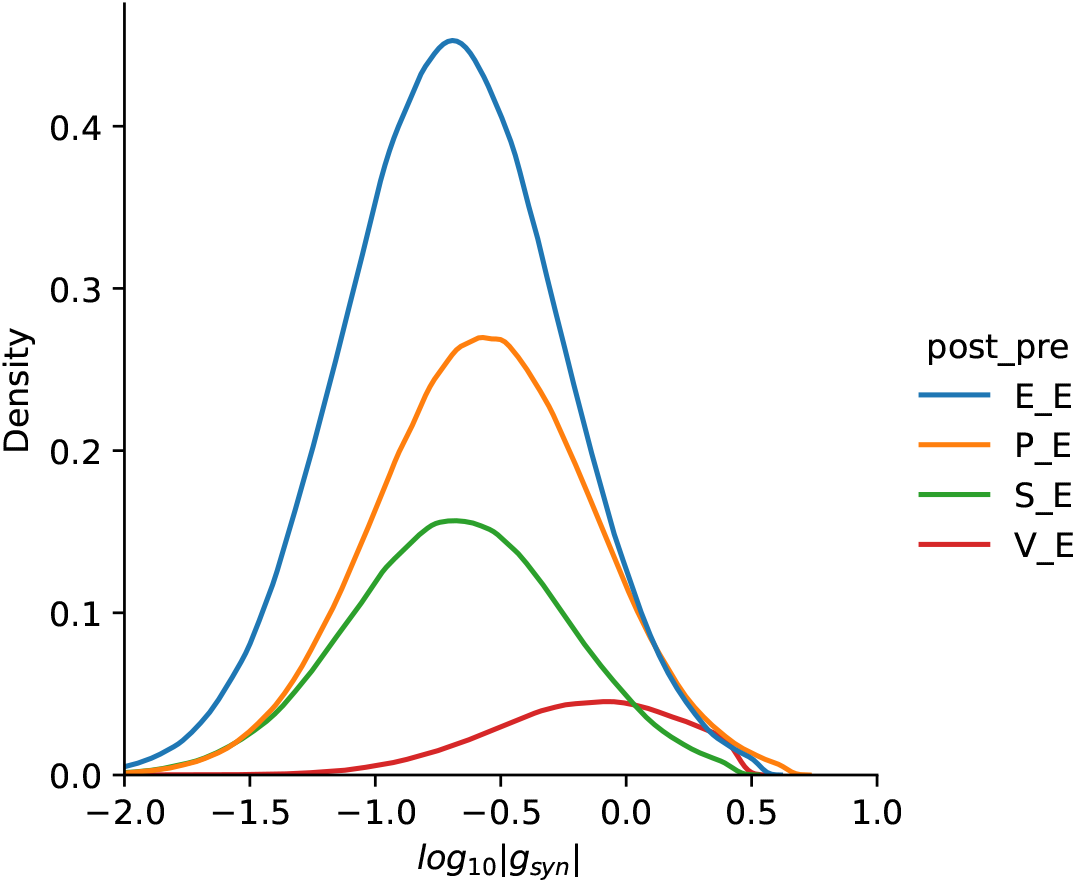
Distribution of non-zero synaptic conductance for E-E, E-P, P-E, P-P connections.

**Table 2:** Network connectivity parameters

| Src \ Tar | Conn. Prob. |  |  |  | Cond. (nS) |  |  |  |
| --- | --- | --- | --- | --- | --- | --- | --- | --- |
|  | PC | PV | SST | VIP | PC | PV | SST | VIP |
| PC | 0.1 | 0.6 | 0.55 | - | 0.20 | 0.08 | 0.16 | - |
| PV | 0.45 | 0.5 | 0.6 | - | 0.27 | 0.46 | 0.45 | - |
| SST | 0.35 | - | 0.5 | 0.5 | 0.21 | - | - | 0.07 |
| VIP | 0.1 | - | 0.45 | 0.6 | 0.78 | - | 0.07 | - |

### Baseline input

To mimic the ongoing activity, each neuron received homogeneous Poisson spike trains. The rates of spike trains to different neuron-type populations were tuned to obtain a baseline firing rate around 2.5 Hz for excitatory and SST neurons, and ≈14 Hz for PV and VIP neurons (spontaneous activity level of free whiskering mice (Yu et al., 2019)). Note the baseline network state was PV-dominated in our model (Supplementary Figure S1 top row).

### Stimulus evoked input

To mimic stimulus-evoked inputs, each population was stimulated by additional inputs with rates 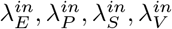 on top of the baseline inputs as shown in Figure 1. Since SST and VIP neurons are mutually coupled, we assumed that modulatory input to VIP neurons is just inverted input to SST neurons as in (Hertäg & Sprekeler, 2019) hence reduced the input dimension to three with 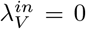. We measured the steady-state response of four populations, 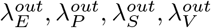 to different levels of modulatory inputs covering a cubic input space. The corresponding output space is restricted due to the interaction between excitation and inhibition as shown in Figure 1. Because the firing rate of the excitatory population was taken as the output, the transfer-function was formulated as 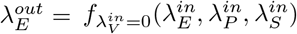 (Figure 1). Negative rates imply a reduction in the input from the baseline input. To obtain a transfer-function with higher resolution, we interpolated the simulated transfer-function with (tri)linear function (Weiser & Zarantonello, 1988) implemented in SciPy library (Virtanen et al., 2020). The analysis of variability transformation was derived from this interpolated neuron transfer-function.

### Trial-by-trial variability of the input

Trial-by-trial variability of the modulatory inputs was modeled as a three-dimensional normal distribution (Figure 3) characterized as:

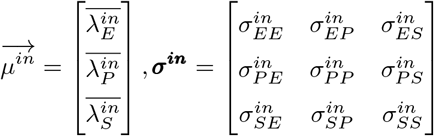

For each setting of mean and covariance matrix, the input rates were sampled (10000-point input cloud) from an arbitrary distribution with given 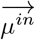 and ***σ^in^*** and corresponding outputs were extrapolated from the neuron transfer-function.

We investigated the transformation of input distribution to output distribution by systematically changing the mean 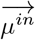, trial-by-trial balance 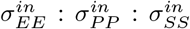, and across trial covariances 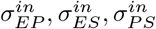 (assuming 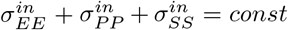). A negative ratio of trial-by-trial input variance (anti-correlated across trials) was generated by flipping the sign of modulatory inputs:

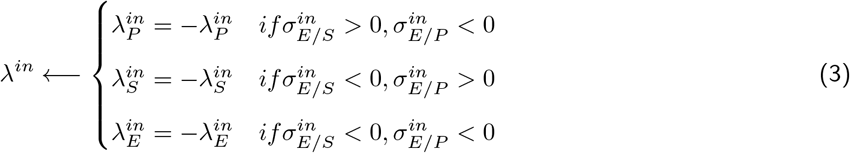

Covariance matrices being not positive semidefinite were discarded. These are marked as white space in the Figure 3 B,C.

To confirm the analysis, we simulated the three network states in Figure 3A right with different covariance matrices. The covariance matrices were chosen to show the trend of variability modulation regarding trial-by-trial covariance and trial-by-trial balance (ratio of input variances): for the former factor, we chose four settings where both pairs, E-P and E-S, had low trial-by-trial input covariance, E-P had large covariance, E-S had large covariance, and both had large covariance (Figure 5 A upper); for the latter factor, we chose four settings where both pairs had low trial-by-trial input variance ratio, both had high ratio, E-S had negative ratio, and E-P had negative ratio (Figure 4 A upper).

For each network state (mean input) and covariance matrix, we simulated 100 trials for 1000 ms each with 250 ms preparation to reach the operating point, 250 ms to reach the stable state after injecting modulatory inputs, and last 500 ms were taken as the steady-state response (Figure 5C and Figure 4C). Firing rate traces of four populations and corresponding spike raster are illustrated in Supplementary Figure S1. The result of trial-by-trial variability control for the PV-dominated regime is shown in Figure 5 and Figure 4, and the result for the SST-dominated regime is given in Supplementary Figure S4.

### Measurement of output trial-by-trial variability

For each simulation, we used the mean firing rate of each population (E, P, and S) in the last 500 ms as the steady-state response rate. Across 100 trials, we calculated the covariance matrix of their steady-state response rate (Figure 5 A bottom and Figure 4 A bottom). Diagonal values in the output covariance matrice denote the trial-by-trial variance of each population and the rest values denote trial-by-trial covariance between populations.

### Simulation and data analysis tools

The simulations were performed using NEST simulator (Jordan et al., 2019). Differential equations were integrated using a fixed timestep of 0.1 ms. The analysis of simulated network activity was done using customized code written in Python. The results were visualized using matplotlib. The code for simulating the network and visual illustrations will be shared online on GitHub.

## Acknowledgments

We thank Dr. Jens Hjerling-Leffler, Dr. Pascal Helson, Emil Waernberg and Movitz Lenninger for helpful discussions. This work was funded in parts by Swedish Research Council (VR), StratNeuro, Digital Futures and Inst. of Advanced Studies (University of Strasbourg, France) Fellowship grants.

---

**Figure S1:**
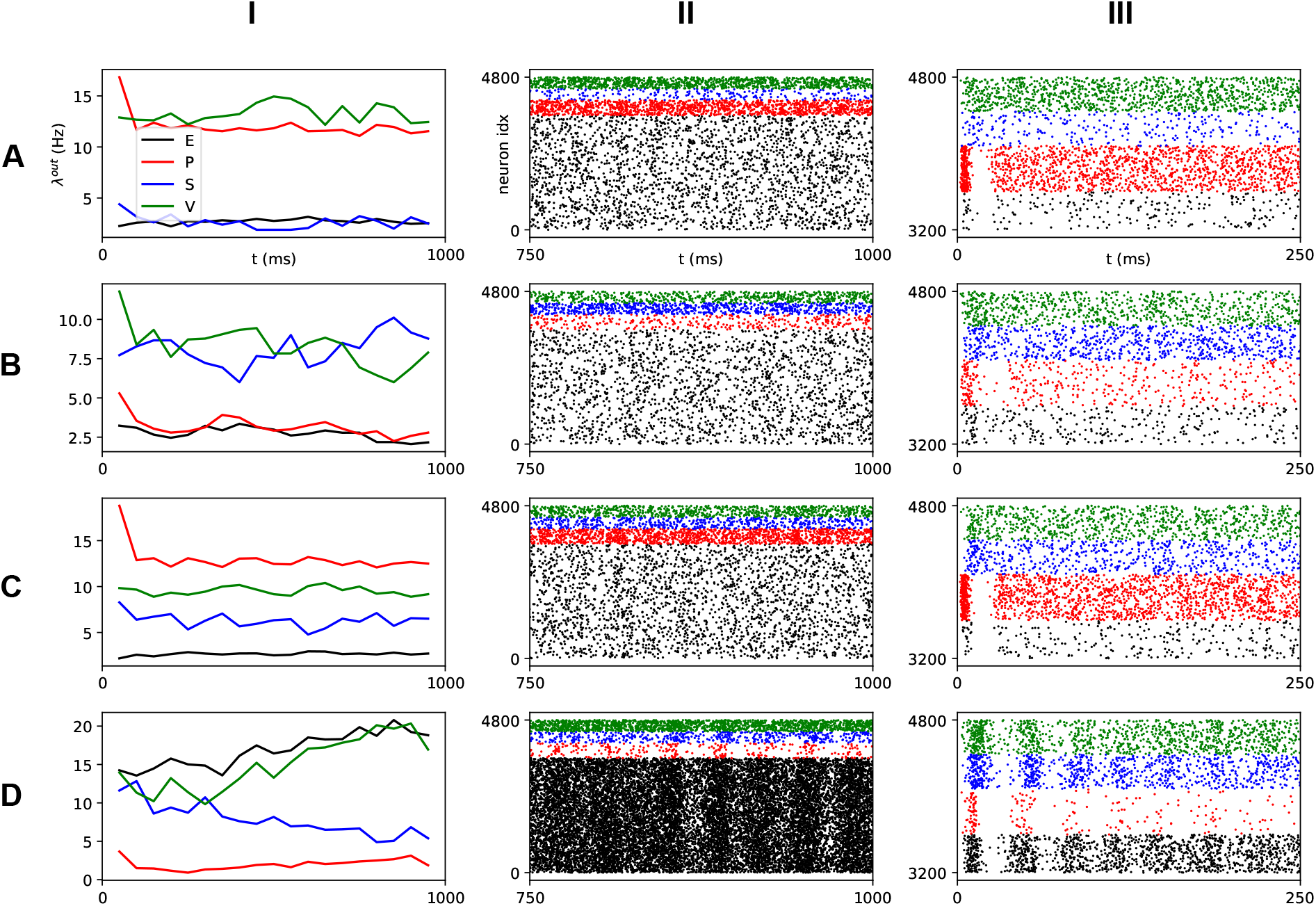
Firing rate and spike raster for a PV-dominated, SST-dominated and both influencing network states. (**A-I**) Average firing rate of four populations in PV-dominated regime (Figure 3 B,C top row) initialized at random state; (**A-II**) Spike aster of all 4800 neurons for last 250ms (colors denote the corresponding population in column **I**); (**A-III**) Zoom-in of initial 250ms for a subset of the whole network. (**B,C**) Same arrangement as in the panel **A** for a SST-dominated regime and a PV-SST driven regime. (**D**) The network can exhibit stochastic oscillation between populations given strong inputs. In our analysis, we avoided such strong inputs and assumed that the network remained in an asynchronous-irregular activity state.

**Figure S2:**
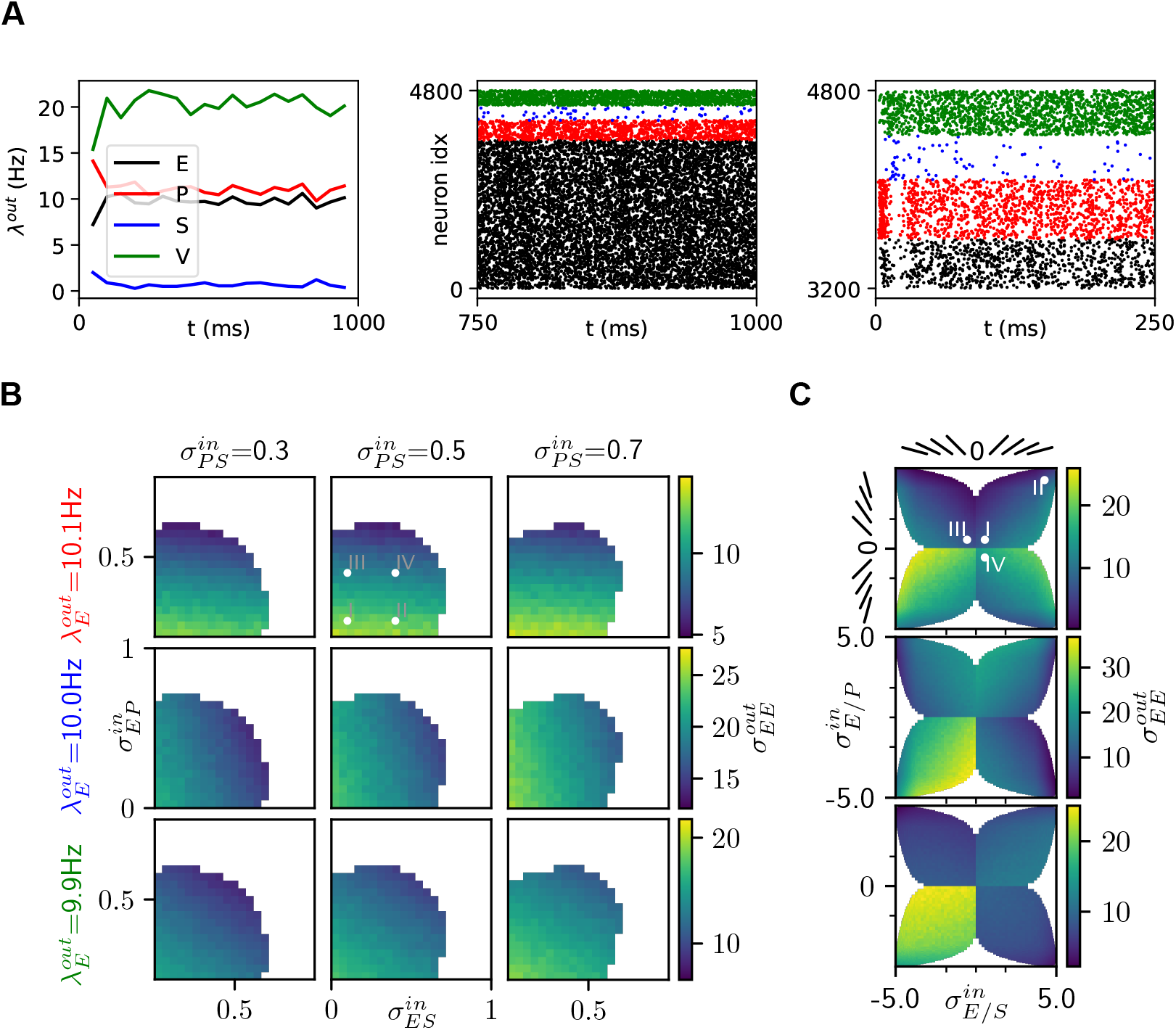
Trial-by-trial variability in EPSV network for high output firing rate (10 Hz) state. (**A** left) Average firing rate, of four populations for 1000ms trial in PV-dominated regime. (**A** middle) Spike raster plot of all 4800 neurons for the last 250ms (colors denote the corresponding population in the left panel); (**A** right) Zoom-in of initial 250ms for a subset of the whole network. (**B**) Trial-by-trial output variability as a function of the trial-by-trial input covariance with fixed variance 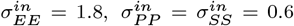 in different network regimes (Top: PV-dominated, Middle: SST-dominated, Bottom: both PV-SST driven). (**C**) Trial-by-trial output variability as a function of the trial-by-trial variance ratio with fixed covariance 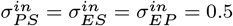. Slanted bars indicate the orientation of the input point cloud. Roman numerals in the panels **B** and **C** refer to the four cases simulated in Figure S3.

**Figure S3:**
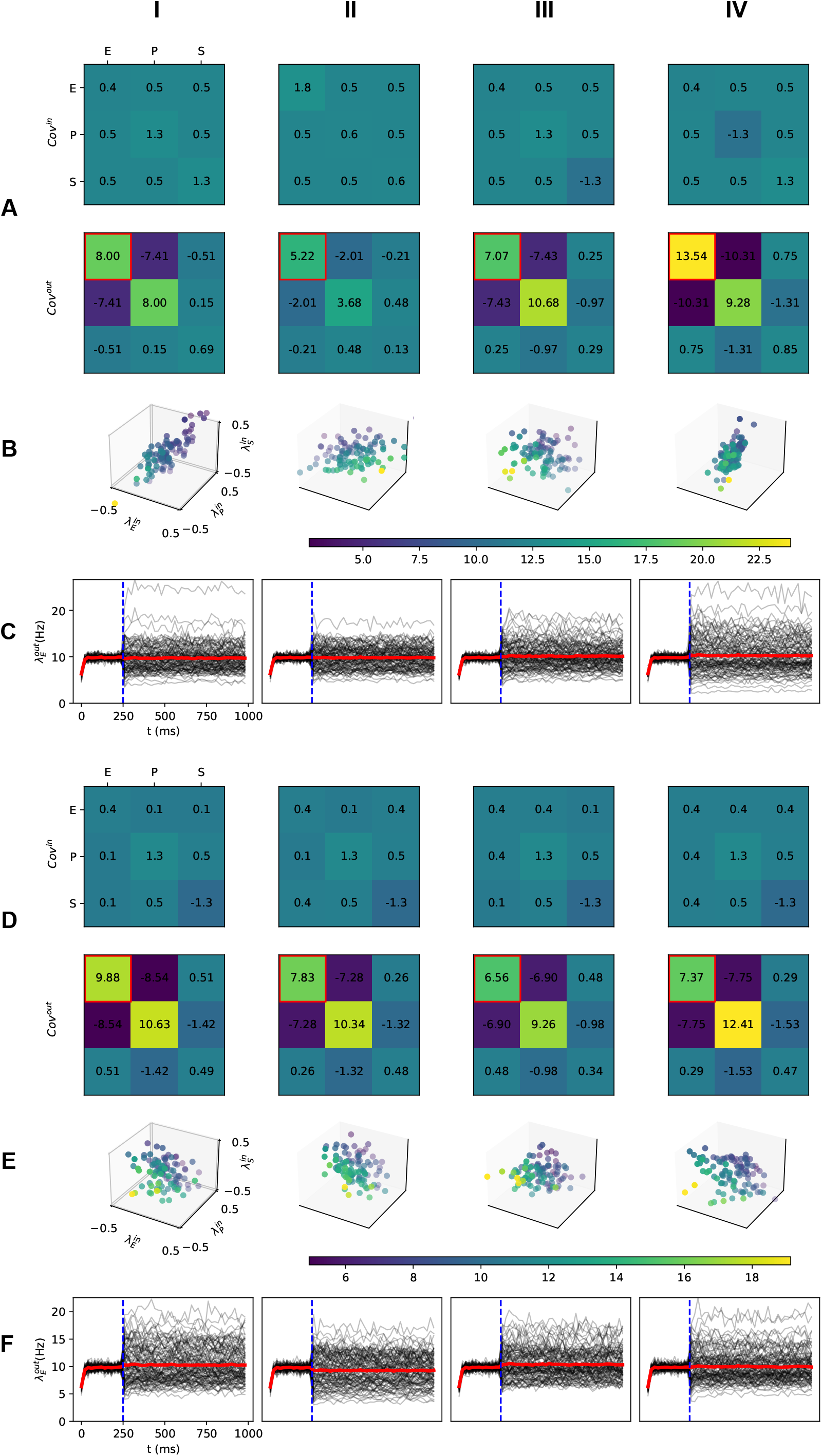
Trial-by-trial variability in a PV-dominated regime for high output firing (10 Hz) state. (**A top**) The input covariance matrix is used to sample inputs across trials (**B**). From **I** to **II**, the trial-by-trial input variance ratio between E and SST/PV populations were increased. In column **III** and **IV**, negative variance in *Cov^in^* denotes the negative slope of input point clouds as shown in Figure 2 F. The network operated in a state corresponding to the middle row in Figure 3 C. (**A bottom**) Output covariance matrix across 100 trials where the input for each trial was sampled from the covariance matrix in **A top**. The output variance of the excitatory population is marked with a red square. (**B**) Trial-by-trial input point clouds sampled from given covariance matrix as columns **I** to **IV** in **A top** (normalized range). **C** PSTH of excitatory population response. Black lines: individual trial. Red line: average response over 100 trials. The stimulus was provided at 250ms, and the output covariance matrix in **A bottom** is calculated for the last 500ms. Note that the stimulus did not involve any change in the input mean, only the input covariance matrix was altered. (**Dtop**) The input covariance matrix for sampling inputs across different trials. From **I** to **II** (**III**), the trial-bytrial input covariance between E and SST (PV) populations were increased. In column **IV**, both covariances were increased. (**D bottom**) Output covariance matrix across 100 trials where the input for each trial was sampled from the covariance matrix in **D top**. The output variance of the excitatory population is marked with a red square. (**E**) trial-by-trial input point clouds sampled from given covariance matrix as columns **I** to **IV** in **D top** (normalized range). (**F**) PSTH of excitatory population response. Black lines: individual trial. Red line: average response over 100 trials. The stimulus was given at 250ms, and the output covariance matrix in **D bottom** is calculated for the last 500ms. The trend is consistent with Figure 3 B,C middle row. Exact values are slightly different due to linear interpolation of neuron transfer-function and limited sampling size for simulations while the actual underlying iso-firing rate surfaces are nonlinear Figure 1 D right.

**Figure S4:**
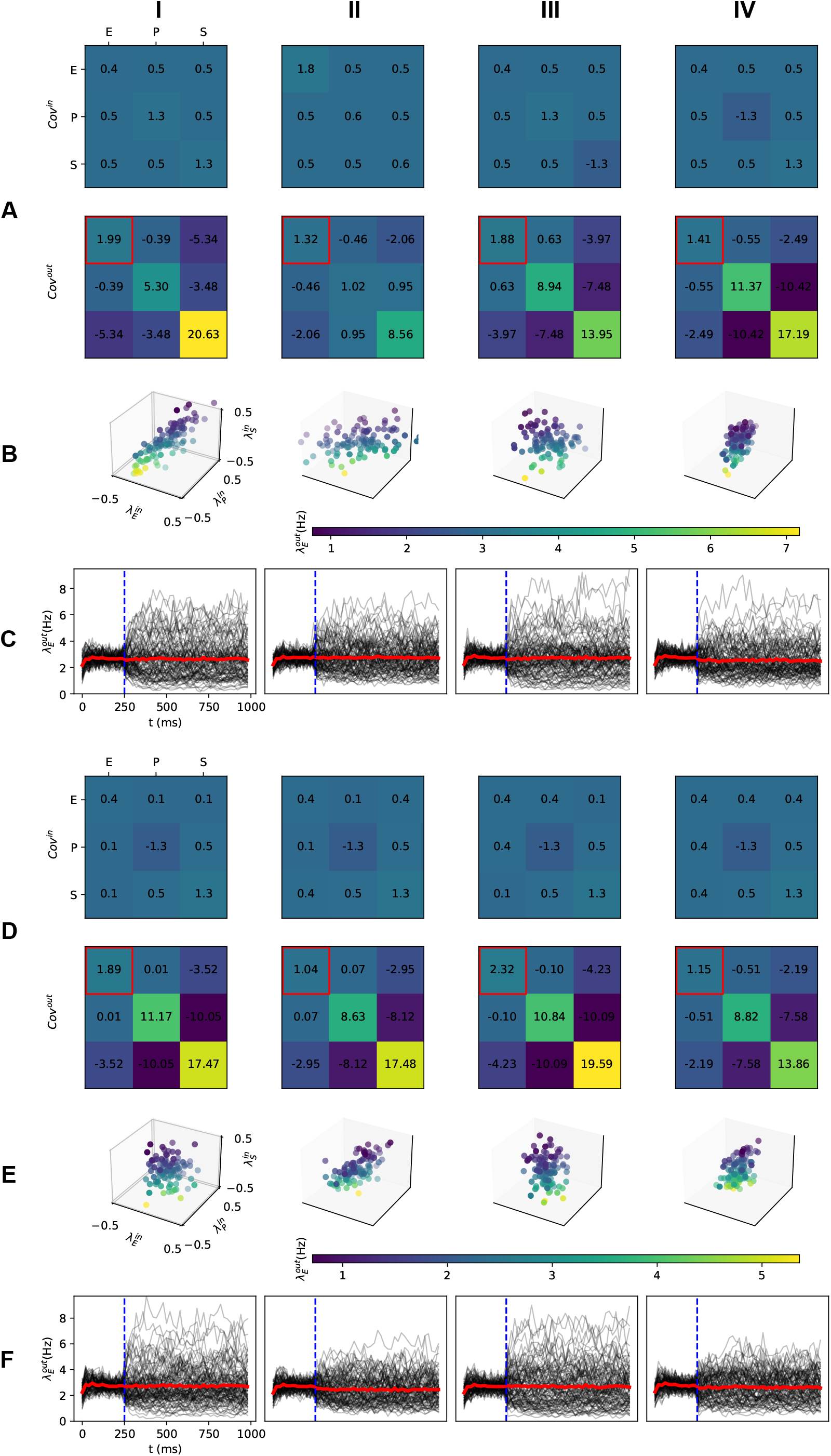
Simulation result of trial-by-trial variability in a SST-dominated regime for low output firing state. (**A top**) The input covariance matrix for sampling inputs for each trial (**B**). From **I** to **II**, the trial-by-trial input variance ratio between E and SST/PV populations were increased. In column **III** and **IV**, negative variance in *Cov^in^* denotes the negative slope of input point clouds as shown in Figure 2 F. The network operates in a state corresponding to the middle row in Figure 3C. (**A bottom**) Output covariance matrix across 100 trials where inputs for each trial were sampled from the covariance matrix in **A top**. The output variance of the excitatory population is marked with a red square. (**B**) Trial-by-trial input point clouds sampled from given covariance matrix as columns **I** to **IV** in **A top** (normalized range). **C** PSTH of excitatory population response. Black lines: individual trial. Red line: average response over 100 trials. Stimulus is given at 250ms, and output covariance matrix in **A bottom** is calculated for the last 500ms. (**Dtop**) The input covariance matrix for sampling input for each trial. From **I** to **II** (**III**), the trial-by-trial input covariance between E and SST (PV) populations were increased. In column **IV**, both covariances were increased. (**D bottom**) Output covariance matrix across 100 trials where inputs for each trial were sampled from the covariance matrix in **D top**. The output variance of the excitatory population is marked with a red square. (**E**) Trial-by-trial input point clouds sampled from given covariance matrix as columns **I** to **IV** in **D top** (normalized range). (**F**) PSTH of excitatory population response. Black lines: individual trial. Red line: average response over 100 trials. Stimulus is given at 250ms, and the output covariance matrix in **D bottom** is calculated for the last 500ms. The trend is consistent with Figure 3 B,C middle row. Exact values are slightly different due to linear interpolation of neuron transfer-function and limited sampling size for simulations while the actual underlying iso-firing rate surfaces are nonlinear Figure 1 D right.

## References

Arazi, A., Censor, N., & Dinstein, I. (2017). Neural variability quenching predicts individual perceptual abilities. Journal of Neuroscience, 37, 97–109.

Arieli, A., Sterkin, A., Grinvald, A., & Aertsen, A. (1996). Dynamics of ongoing activity: explanation of the large variability in evoked cortical responses. Science, 273, 1868–1871.

Brette, R., & Gerstner, W. (2005). Adaptive exponential integrate-and-fire model as an effective description of neuronal activity. Journal of Neurophysiology, 94, 3637–3642.

Brunel, N. (2000). Dynamics of sparsely connected networks of excitatory and inhibitory spiking neurons. Journal of computational neuroscience, 8, 183–208.

Bujan, A. F., Aertsen, A., & Kumar, A. (2015). Role of input correlations in shaping the variability and noise correlations of evoked activity in the neocortex. Journal of Neuroscience, 35, 8611–8625.

Churchland, M. M., Byron, M. Y., Cunningham, J. P., Sugrue, L. P., Cohen, M. R., Corrado, G. S., Newsome, W. T., Clark, A. M., Hosseini, P., Scott, B. B. et al. (2010). Stimulus onset quenches neural variability: a widespread cortical phenomenon. Nature Neuroscience, 13, 369–378.

De Luna, P., Veit, J., & Rainer, G. (2017). Basal forebrain activation enhances between-trial reliability of low-frequency local field potentials (lfp) and spiking activity in tree shrew primary visual cortex (v1). Brain Structure and Function, 222, 4239–4252.

Deco, G., & Hugues, E. (2012). Neural network mechanisms underlying stimulus driven variability reduction. PLoS Computational Biology, 8, e1002395.

Doiron, B., Litwin-Kumar, A., Rosenbaum, R., Ocker, G. K., & Josić, K. (2016). The mechanics of state-dependent neural correlations. Nature Neuroscience, 19, 383–393.

Hahn, G., Bujan, A. F., Frégnac, Y., Aertsen, A., & Kumar, A. (2014). Communication through resonance in spiking neuronal networks. PLoS Computational Biology, 10, e1003811.

Hahn, G., Kumar, A., Schmidt, H., Knösche, T. R., & Deco, G. (2022). Rate and oscillatory switching dynamics of a multilayer visual microcircuit model. Elife, 11, e77594.

Hertäg, L., & Sprekeler, H. (2019). Amplifying the redistribution of somato-dendritic inhibition by the interplay of three interneuron types. PLoS Computational Biology, 15, e1006999.

Isaacson, J. S., & Scanziani, M. (2011). How inhibition shapes cortical activity. Neuron, 72, 231–243.

Jiang, X., Shen, S., Cadwell, C. R., Berens, P., Sinz, F., Ecker, A. S., Patel, S., & Tolias, A. S. (2015). Principles of connectivity among morphologically defined cell types in adult neocortex. Science, 350, aac9462.

Jordan, J., Deepu, R., Mitchell, J., Eppler, J. M., Spreizer, S., Hahne, J., Thomson, E., Kitayama, I., Peyser, A., Fardet, T. et al. (2019). NEST 2.18. 0. Technical Report Jülich Supercomputing Center.

Kanashiro, T., Ocker, G. K., Cohen, M. R., & Doiron, B. (2017). Attentional modulation of neuronal variability in circuit models of cortex. Elife, 6, e23978.

Kepecs, A., & Fishell, G. (2014). Interneuron cell types: fit to form and formed to fit. Nature, 505, 318.

Kremkow, J., Aertsen, A., & Kumar, A. (2010). Gating of signal propagation in spiking neural networks by balanced and correlated excitation and inhibition. Journal of neuroscience, 30, 15760–15768.

Kuhn, A., Aertsen, A., & Rotter, S. (2004). Neuronal integration of synaptic input in the fluctuation-driven regime. Journal of Neuroscience, 24, 2345–2356.

Larkum, M. (2013). A cellular mechanism for cortical associations: an organizing principle for the cerebral cortex. Trends in Neurosciences, 36, 141–151.

Lee, B., Shin, D., Gross, S. P., & Cho, K.-H. (2018). Combined positive and negative feedback allows modulation of neuronal oscillation frequency during sensory processing. Cell Reports, 25, 1548–1560.

Litwin-Kumar, A., & Doiron, B. (2012). Slow dynamics and high variability in balanced cortical networks with clustered connections. Nature Neuroscience, 15, 1498–1505.

Marshel, J. H., Kim, Y. S., Machado, T. A., Quirin, S., Benson, B., Kadmon, J., Raja, C., Chibukhchyan, A., Ramakrishnan, C., Inoue, M. et al. (2019). Cortical layer–specific critical dynamics triggering perception. Science, 365, eaaw5202.

Naka, A., Veit, J., Shababo, B., Chance, R. K., Risso, D., Stafford, D., Snyder, B., Egladyous, A., Chu, D., Sridharan, S. et al. (2019). Complementary networks of cortical somatostatin interneurons enforce layer specific control. Elife, 8, e43696.

Oram, M. W. (2011). Visual stimulation decorrelates neuronal activity. Journal of Neurophysiology, 105, 942–957.

Renart, A., De La Rocha, J., Bartho, P., Hollender, L., Parga, N., Reyes, A., & Harris, K. D. (2010). The asynchronous state in cortical circuits. Science, 327, 587–590.

Rowland, J. M., van der Plas, T. L., Loidolt, M., Lees, R. M., Keeling, J., Dehning, J., Akam, T., Priesemann, V., & Packer, A. M. (2021). Perception and propagation of activity through the cortical hierarchy is determined by neural variability. bioRxiv,.

Rudy, B., Fishell, G., Lee, S., & Hjerling-Leffler, J. (2011). Three groups of interneurons account for nearly 100% of neocortical gabaergic neurons. Developmental Neurobiology, 71, 45–61.

Sanzeni, A., Akitake, B., Goldbach, H. C., Leedy, C. E., Brunel, N., & Histed, M. H. (2020). Inhibition stabilization is a widespread property of cortical networks. Elife, 9, e54875.

Schnepel, P., Kumar, A., Zohar, M., Aertsen, A., & Boucsein, C. (2015). Physiology and impact of horizontal connections in rat neocortex. Cerebral Cortex, 25, 3818–3835.

Silver, R. A. (2010). Neuronal arithmetic. Nature Reviews Neuroscience, 11, 474–489.

Tetzlaff, T., Helias, M., Einevoll, G. T., & Diesmann, M. (2012). Decorrelation of neural-network activity by inhibitory feedback. PLoS Computational Biology, 8.

Veit, J., Hakim, R., Jadi, M. P., Sejnowski, T. J., & Adesnik, H. (2017). Cortical gamma band synchronization through somatostatin interneurons. Nature Neuroscience, 20, 951–959.

Virtanen, P., Gommers, R., Oliphant, T. E., Haberland, M., Reddy, T., Cournapeau, D., Burovski, E., Peterson, P., Weckesser, W., Bright, J., van der Walt, S. J., Brett, M., Wilson, J., Millman, K. J., Mayorov, N., Nelson, A. R. J., Jones, E., Kern, R., Larson, E., Carey, C. J., Polat, İ., Feng, Y., Moore, E. W., VanderPlas, J., Laxalde, D., Perktold, J., Cimrman, R., Henriksen, I., Quintero, E. A., Harris, C. R., Archibald, A. M., Ribeiro, A. H., Pedregosa, F., van Mulbregt, P., & SciPy 1.0 Contributors (2020). SciPy 1.0: Fundamental Algorithms for Scientific Computing in Python. Nature Methods, 17, 261–272.

Waschke, L., Kloosterman, N. A., Obleser, J., & Garrett, D. D. (2021). Behavior needs neural variability. Neuron, 109, 751–766.

Weiser, A., & Zarantonello, S. E. (1988). A note on piecewise linear and multilinear table interpolation in many dimensions. Mathematics of Computation, 50, 189–196.

White, B., Abbott, L. F., & Fiser, J. (2012). Suppression of cortical neural variability is stimulus-and state-dependent. Journal of Neurophysiology, 108, 2383–2392.

Yu, J., Hu, H., Agmon, A., & Svoboda, K. (2019). Recruitment of gabaergic interneurons in the barrel cortex during active tactile behavior. Neuron, 104, 412–427.

